# Preclinical exploration of the DNA Damage Response pathway using the interactive neuroblastoma cell line explorer CLEAN

**DOI:** 10.1101/2023.09.17.557904

**Authors:** Jonatan L. Gabre, Peter Merseburger, Arne Claeys, Joachim Siaw, Sarah-Lee Bekaert, Frank Speleman, Bengt Hallberg, Ruth H. Palmer, Jimmy Van den Eynden

## Abstract

Neuroblastoma (NB) is the most common cancer in infancy with an urgent need for more efficient targeted therapies. The development of novel (combinatorial) treatment strategies relies on extensive explorations of signaling perturbations in neuroblastoma cell lines, using RNA-Seq or other high throughput technologies (e.g., phosphoproteomics). This typically requires dedicated bioinformatics support, which is not always available. Additionally, while data from published studies are highly valuable and raw data (e.g., fastq files) are nowadays released in public repositories, data processing is time-consuming and again difficult without bioinformatics support. To facilitate NB research, more user-friendly and immediately accessible platforms are needed to explore newly generated as well as existing high throughput data. To make this possible, we developed an interactive data centralization and visualization web application, called CLEAN (the Cell Line Explorer web Application of Neuroblastoma data; https://ccgg.ugent.be/shiny/clean/). By focusing on the regulation of the DNA damage response, a therapeutic target of major interest in neuroblastoma, we demonstrate how CLEAN can be used to gain novel mechanistic insights and identify putative drug targets in neuroblastoma.

## INTRODUCTION

Neuroblastoma (NB) is the most common cancer in infancy and accounts for approximately 15% of pediatric cancer-related deaths (1). Despite initial positive treatment response, high-risk NB frequently leads to relapse, resulting in a survival rate as low as 35% (2). Our groups and others have recently demonstrated in different preclinical NB animal models that targeting the DNA damage response (DDR) pathway is a promising treatment strategy for high-risk NB (3–6).

Successful investigations of new or combinatorial drug targets are often based on extensive *in vitro* experiments in cell lines, followed by high throughput experiments such as RNA-Seq and/or phosphoproteomics (3,7,8). Over the past few decades, generation of these types of data has become increasingly efficient, leading to a strong reduction in price as well as increase in accuracy. This has led to a surge in omics data abundance in the NB, as well as most other pre-clinical fields (9) – a development that has brought both opportunities and challenges (10). While each omics study generates tens of thousands of data points, only a fraction of this information is used to draw conclusions and an even smaller proportion is addressed in the final manuscript of any study. A single perturbation, even in a relatively homogenous system like immortal cell lines, causes major, complex downstream effects at several regulatory levels (3). A complete understanding of the cellular response is a task that goes beyond the scope of a single study. Providing real access to these datasets would let us deepen our understanding of each experiment and move the entire field of NB research forward substantially. Despite attempts to increase reproducibility, by mandating upload of raw data to online repositories and encouraging code sharing (11–13), proper accessibility to this excess information remains a problem.

To meet the need for more user-friendly, simpler and faster ways to explore and access this goldmine of experimental data, we developed a highly interactive data centralization and visualization web application, called CLEAN (the

Cell Line Explorer web Application of Neuroblastoma data; https://ccgg.ugent.be/shiny/CLEAN/). We illustrate the CLEAN functionality and its added value for translational NB research by providing new mechanistic insights in the DDR regulation and identifying novel therapeutic targets for future studies.

## RESULTS

### CLEAN is an explorable and standardized neuroblastoma cell line data repository

The aim of the CLEAN initiative is to make all experimental NB cell line data that involve RNA-Seq and/or phosphoproteomics experiments publicly available in an easy-to-use and interactive format. We selected all available studies from the Gene Expression Omnibus (GEO) (14) and the ProteomeXchange (PRX) (15) that fulfilled the following inclusion criteria: 1) NB cell lines; 2) minimally 2 replicates per condition and 3) some kind of experimental perturbation (e.g., gene knockout, drug treatment; see Methods for full criteria and data processing). All data were reprocessed using standardized pipelines, resulting in the availability of 66 RNA-Seq and 8 phosphoproteomics studies, with 6 chemotherapeutic drug treatments (chemo), 29 compound treatments, 49 drug inhibitions, 40 gene knockdowns (KD), 15 gene knockouts (KO) and 27 gene overexpressions (OE; Fig. 1).

**Figure 1.**
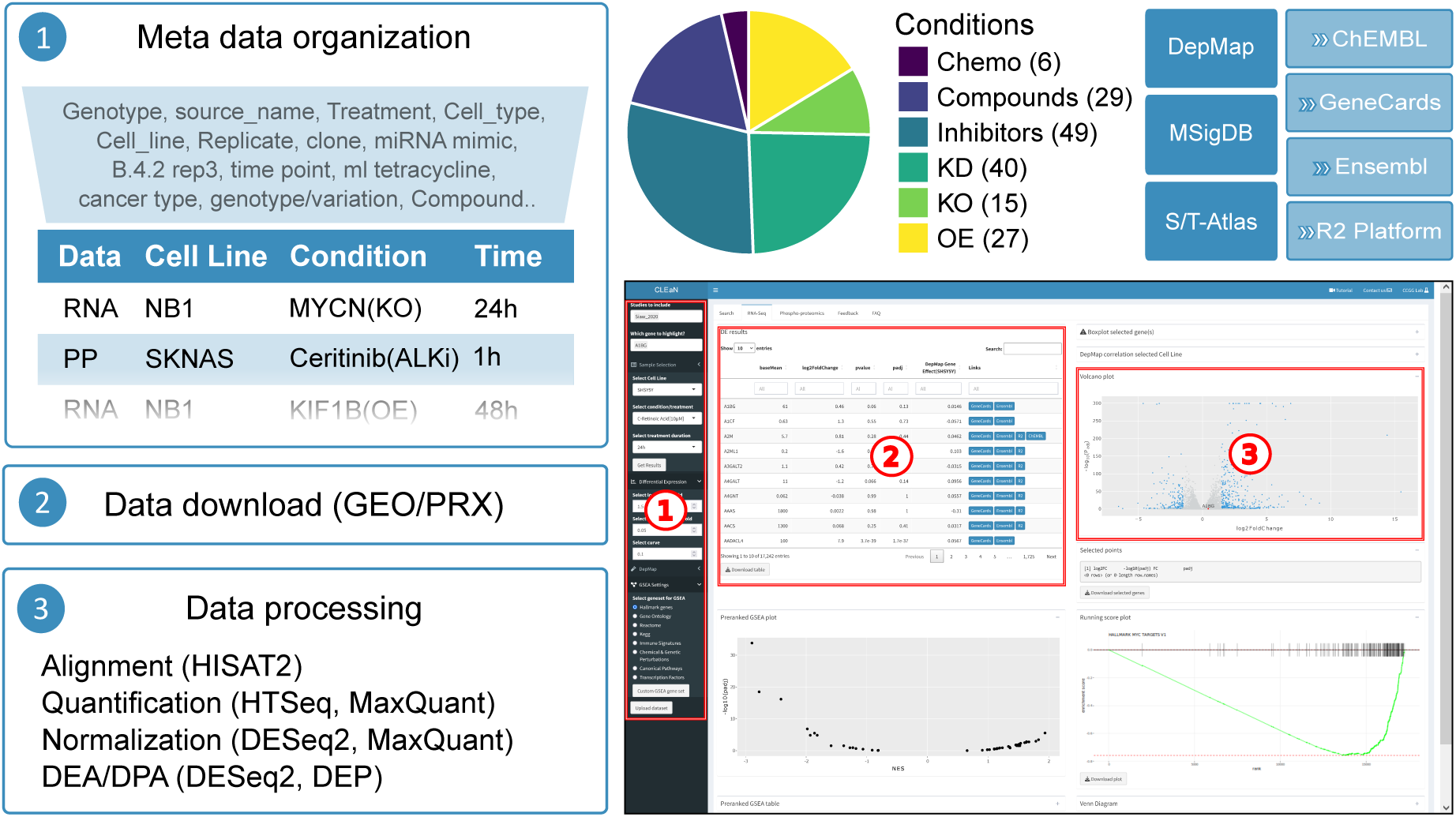
Overview of data processing and availability in CLEAN. Overview of default data processing in CLEAN (left). Pie chart showing the different experimental perturbations that are currently implemented in CLEAN with precise number of studies indicated between parentheses (top-mid). CLEAN screenshot of the default RNA-Seq tab, indicating the 3 main components: (1) the sidebar takes user input and updates what’s in the main panel; (2) a searchable table with detailed differential gene expression or related information and (3) an interactive plot (bottom-right). External data sources that are directly (left) or indirectly (via hyperlinks) (top-right), integrated in CLEAN.

### Interactively exploring high throughput data derived from neuroblastoma cell lines using CLEAN

The CLEAN web application contains 3 main tabs: a *search* tab and 2 data tabs (i.e., *RNA-Seq* and *Phosphoproteomics*). Each tab contains a side bar on the left that accepts user input to query studies and data and a main panel on the right containing a series of downloadable tables and related interactive plots (Fig. 1).

The search tab can be used to quickly check 1) whether RNA-Seq or phosphoproteomics data are available for a specified cell line and/or perturbation/treatment of interest (using a studies overview table in the main panel); 2) which previous studies contain significantly up- or downregulated genes/phosphosites, based on user-defined search criteria (i.e., log2 foldchange (FC) and *P* value*; Sidebar: Search for DE/DP Genes/PP-Sites*); 3) which previous studies contain specific enrichments of user defined gene sets (*Sidebar: GSEA*).

The 2 data tabs have identical features and functionality. They all take user-specified inputs, either indirectly from the *search* tab or directly after selecting the data of interest in the sidebar (i.e., which studies, cell line, condition, time point and gene(s) of interest). Based on these inputs, standardized differential gene/protein expression information is provided for all studies (i.e., log2FC and *P* values), both in an easily searchable data table format and an interactive volcano plot. The data table contains gene-specific links to complementary resources, such as *GeneCards* (16), *ChEMBL* (17), *Ensembl* (18) and the *R2: Genomics Analysis and Visualization Platform* (http://r2.amc.nl). Relatedly, all findings can be easily correlated to CRISPR/cas9 generated gene effect scores from the *DepMap* database (19), potentiating the identification of essential genes affected by the selected perturbation easy (Fig. 1).

### CLEAN provides different forms of existing and user defined gene set enrichment functionality

CLEAN also contains extensive gene set enrichment analysis (GSEA) functionalities, both in the form of a preranked GSEA and an overrepresentation analysis. The former first ranks the genes based on their *P* values and considers the type of differential expression (up- or downregulated) and the enrichment results (i.e., normalized enrichment scores and *P* values) are then visualized in an interactive volcano plot at the dataset level, dynamically linked with a running score plot at the individual gene set level. The latter, overrepresentation analysis is performed using Fisher’s exact test, considering a set of genes that are defined based on user-determined cut-off criteria for differential expression. Enrichment results (i.e., odds ratios and *P* values) are again visualized in tables and volcano plots. CLEAN also allows the direct cross-study and -condition comparisons of up to 4 different studies using interactive Venn diagram, allowing for quick enrichment analyses on intersecting or unique gene sets.

The most common gene sets from the Molecular Signatures Database (MSigDB) are available (20) and a user also has the option to upload any custom gene set of interest. For phosphoproteomics data analysis, a protein kinase data set from the recently published atlas of serine/threonine substrate specificities is provided (21).

### Searching the CLEAN repository for previous experiments that altered the DDR pathway in neuroblastoma cell lines

Given the emerging evidence of a therapeutically targetable role of the DDR in NB (e.g., using AURKA, ATR or CHK1 inhibitors) (3,5,22), we used the CLEAN *search tab* (available at the home page) to identify other known drugs that inhibit the DDR and could be used as putative combinatorial treatments.

We employed a curated set of 276 DDR-related genes derived from *Knijnenburg et al.* (23) and uploaded these genes as a custom gene set in CLEAN (*Search tab, Sidebar: GSEA*). Querying the CLEAN repository for enrichments of these DDR genes in significantly downregulated genes (default settings: log2FC < − 1.5 at 5% FDR) resulted in several RNA-Seq studies with significant (P_adj_ < 0.05) enrichments. As expected, DDR enrichments were found after treating NB cells with the ATR inhibitor elimusertib (CLB-GE cells, 48h treatment; *P*_adj_ = 3.1e-14). Interestingly and in line with our recent experimental findings (6), strong enrichments were also observed after treating NB cells for 24h with the ALK inhibitor lorlatinib (NB1 cells; *P*_adj_ = 7.1e-14), for 48h with the PI3K/mTOR inhibitor dactolisib (NB1 cells; *P*_adj_ = 1.2e-13) or for 24h with the MDM2 inhibitor idasanutlin (NB1691 cells; *P*_adj_ = 1.1e-30). Background information on all these studies is available in CLEAN and after selecting the latter study (24) for further exploration, CLEAN redirects the user to the RNA-Seq tab.

A more detailed differential gene expression analysis of the selected study in the RNA-Seq tab indeed confirms a strong DDR signature (*P* = 1.1e-25), characterized by a downregulation of *RRM1/2*, *EXO1*, *BRCA1*, *FANCI*, *BRIP1*, *CDC25A, POLQ, CHEK1* and many other genes (Fig. 2A-B, Supplementary file 1). We then used the Venn diagram functionality in CLEAN to determine where the downstream signaling pathways upon ATR inhibition (elimusertib), ALK inhibition (lorlatinib), PI3K inhibition (dactolisib) and MDM2 inhibition (idasanutlin) converge. We found 109 commonly downregulated genes that were strongly enriched for the DDR (as expected) and many cell cycle-related processes such as G2M checkpoints and E2F targets, as can be observed when selecting the Canonical Pathways as a data resource for the GSEA (Fig. 3C-D). While some of these genes (e.g., *RRM2, AURKB, CDK2*) have previously been shown to have potential as a drug target in neuroblastoma (22,25,26), many of them are currently underexplored and could serve as an interesting source of further research.

**Figure 2.**
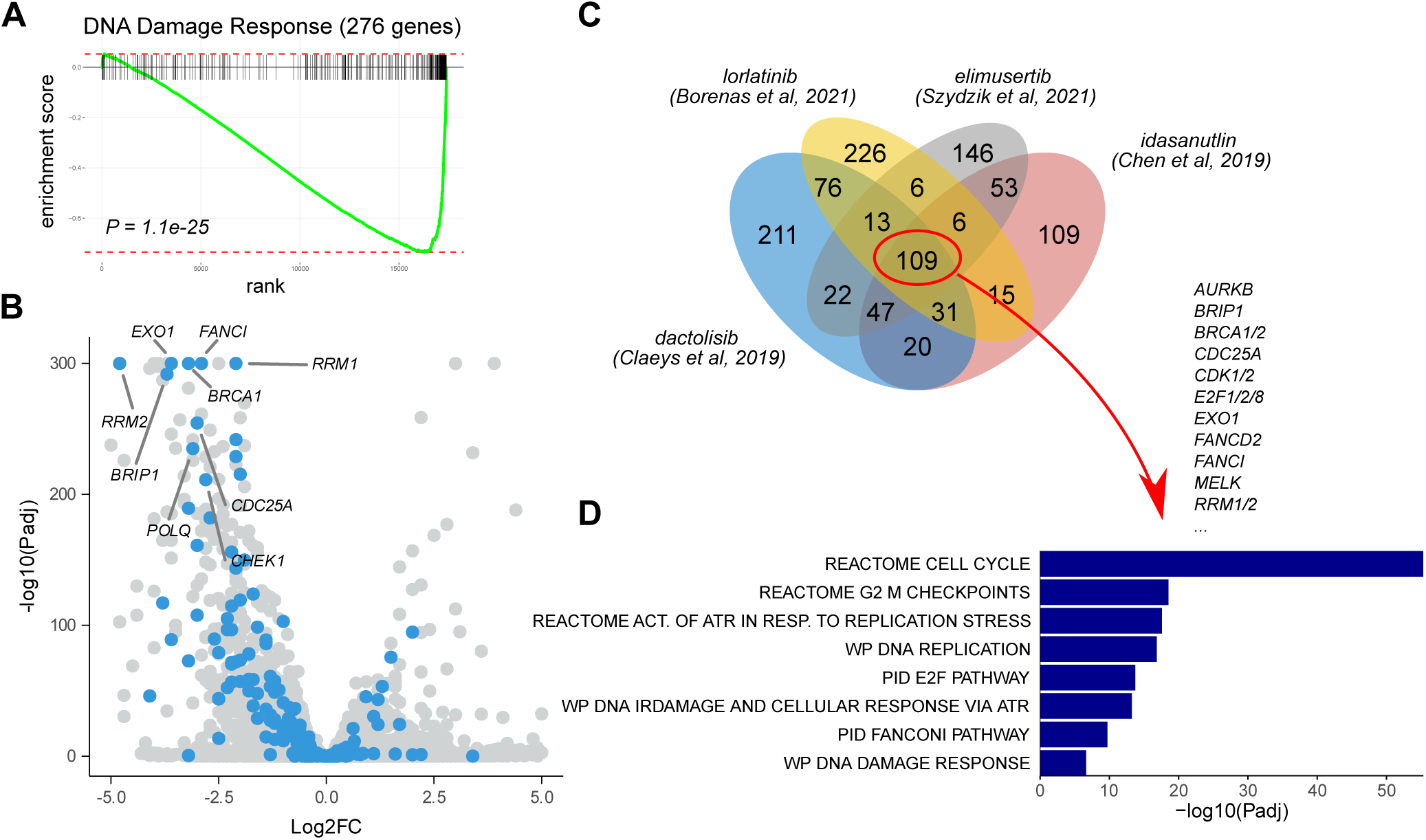
CLEAN-based identification of previously studied drugs that result in a reduced DNA Damage Response transcriptomic signature in neuroblastoma. CLEAN was used to search for previous studies that resulted in significantly downregulated genes (default settings; i.e., logFC < −1.5 at 5% FDR) that were enriched (at 5% FDR) for a custom DNA Damage Response (DDR) gene set (276 genes, as described by *Knijnenburg et al.* (23)). Four retrieved studies were then further explored in the RNA-Seq tab. Running score **(A)** and volcano plot **(B)** for the DDR gene set in the study *of Chen et al, 2019* (*24*). DDR genes indicated in blue in the volcano plot with labelling of genes that are discussed in the main text. **(C)** Venn diagram showing the number of unique and intersecting genes between the 4 selected studies. A GSEA using canonical pathways was then performed in the 109 common genes. Bar plots indicate −log10(P_adj_) values for a selection of enriched pathways as indicated. All plots were generated based on downloaded data from CLEAN. See supplementary file 1 for an illustration of this use case.

**Figure 3.**
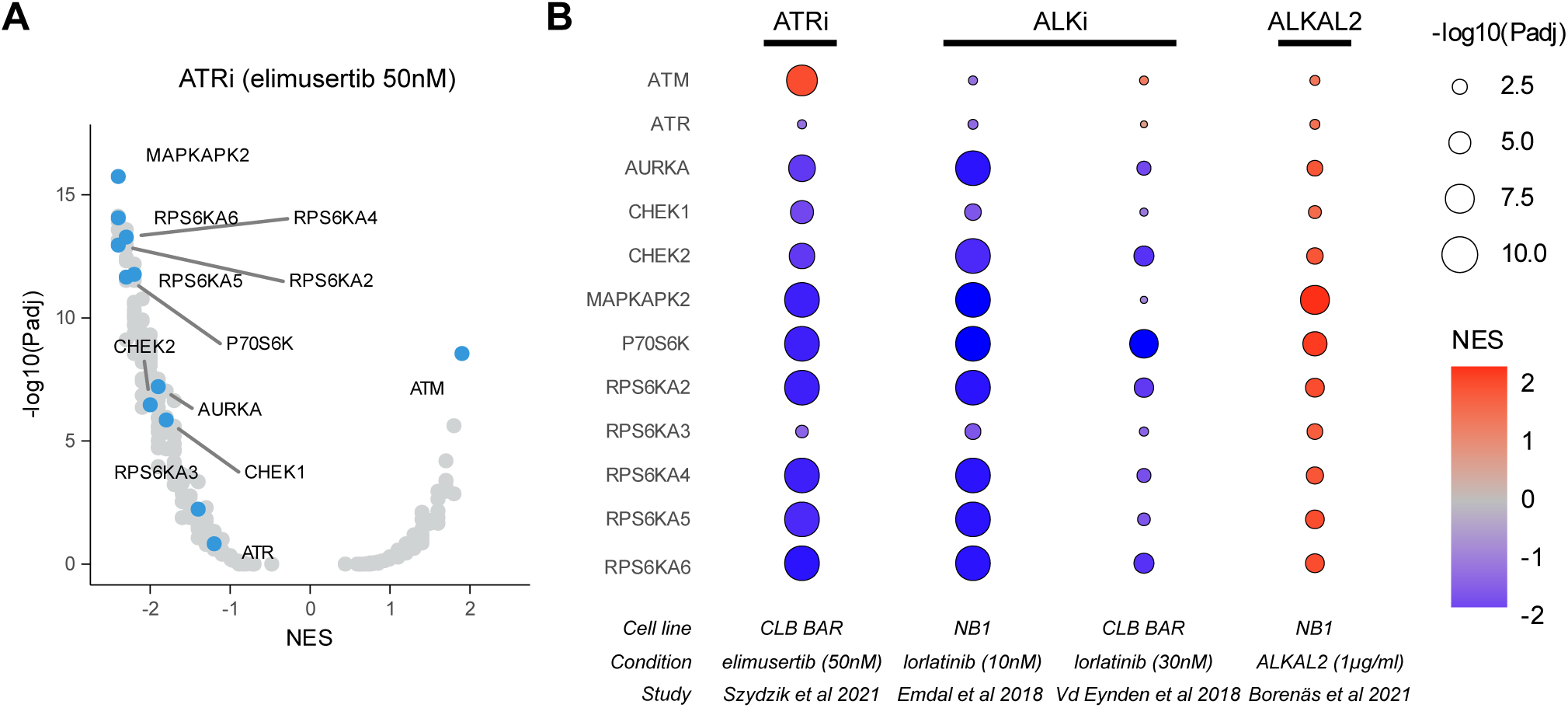
Comparison of phosphomotif-based protein kinase enrichments after ATR and ALK inhibition and/or stimulation. Four studies were selected in CLEAN that had available phosphoproteomics data after drug treatment with elimusertib, lorlatinib or the ALK ligand ALKAL2. The preranked GSEA functionality was then used to search for protein kinase enrichments and results were downloaded. **(A)** Volcano plot showing protein kinase enrichment results after elimusertib treatment in the study of *Szydzik et al, 2021* (3). Protein kinases that are discussed in the main text are labelled. **(B)** Heatmap showing normalized enrichment scores (color scale) and *P* values (size) of the 4 selected studies as indicated on the bottom. See supplementary file 2 for an illustration of this use case. NES, normalized enrichment scores.

### Motif enrichment analyses on previously published phosphoproteomics studies suggest a role for Ribosomal S6 kinases in mediating ALK and ATR inhibition

Apart from the large amount of RNA-Seq data, CLEAN contains phosphoproteomics data that have been previously generated upon drug treatment of NB cell lines (accessible via the *phosphoproteomics tab*). In contrast to RNA-Seq, phosphoproteomics data provide information on earlier and more upstream signal transduction pathways in response to different stimuli. Using this technology, we previously demonstrated that elimusertib-induced ATR inhibition results in an expected dephosphorylation of ATR targets and an increased, compensatory phosphorylation response of ATM targets (3), in keeping with the observations of others (27). Like the RNA-Seq functionality, these data are easily explorable using available interactive tables and volcano plots in CLEAN.

To maximize the efficacy of CLEAN in integrating phosphoproteomics datasets we have implemented a feature based on the recently published atlas of substrate specificities for the human serine/threonine kinome (*Sidebar: GSEA Settings – S/T-Kinases*) (21). This feature allows the analysis of protein kinase enrichments based on differentially phosphorylated phosphosite motifs within the proteins of interest. When performing this analysis on elimusertib-treated NB cells (3), we could indeed confirm a (non-significant) decreased activity of ATR (NES = − 1.2, *P*_adj_ = 0.12) and a strong significant increase in ATM activity (NES = 2, *P*_adj_ = 4.1e-9) upon treatment (Fig. 3A). Additionally, this feature identified decreased activity of the related kinases CHEK1, CHEK2 and AURKA. Given the higher described DDR inhibition upon treatment with lorlatinib, we then hypothesized that lorlatinib would result in similar protein kinase responses. Using the search tab (*Studies* table *– Condition column, search: Lorlatinib*), CLEAN was able to identify two studies that previously explored the phosphoproteomic response to lorlatinib treatment: *Emdal et al., 2018* (NB1 cells) (28) and *Van den Eynden et al., 2018* (CLB-BAR cells) (29). While a motif enrichment analysis did not indicate any significant enrichment for ATM and ATR itself in any of those studies, reduced protein kinase activity was predicted in both studies for the ATR target CHEK1, in agreement with our recent wet-lab experimental findings (6), as well as the ATM target CHEK2 and AURKA (Fig. 3B). Strikingly, mirrored responses were observed upon stimulation of NB1 cells with the ALK ligand ALKAL2 (study of *Borenäs et al., 2021* (*30*)), strongly suggesting that these responses are ALK-specific (Fig. 3B, Supplementary file 2).

The strongest reductions in predicted protein kinase activity upon ATR or ALK inhibition were found for several members of the Ribosomal S6 kinases: RPS6KA2, RPS6KA4, RPS6KA6 (also known as RSK4), P90RSK (also known as MAPKAPK1), MAPKAPK2 and P70S6K (Fig. 3A-B, Supplementary file 2). Several of these kinases are activated by the MAPK/ERK pathway (31), and phosphorylate the ribosome protein S6 (RPS6) (32). Further exploration in CLEAN indicates that this protein is indeed strongly dephosphorylated upon lorlatinib treatment, mainly at S235 (*P*_adj_ = 4.2e-05; log2FC = −1.3). Strikingly, this protein is also highly essential for NB cell lines, as can be observed from the very low DepMap gene effect scores (gene effect score in NB1 cells = −2.47). Interestingly, using the links provided to the *R2: Genomics Analysis and Visualization Platform*, it can be easily shown that high *RPS6KA6* expression is strongly correlated with worse survival in neuroblastoma patients (*P* = 2.07e-12).

## DISCUSSION

CLEAN aims to centralize all available RNA-Seq and phosphoproteomics data derived from NB cell line experiments in a standardized and easily accessible format. It is an ongoing open data science initiative that was developed by a multidisciplinary group of scientists with experience in computational biology, NB and signal transduction. We are planning regular updates (every 3 months) of the available cell line and related gene set data. Additionally, extensions are foreseen for data generated from organoids or NB model organisms (e.g., mouse models). New data sets will be selected from literature or at a user’s request (a contact tab is included in the application). While the CLEAN scope is restricted to NB, its functionality can also be used for other (cancer) cell lines. To facilitate this broader applicability, an option is provided for users to upload other (unpublished) DE analysis results and use all the features provided by CLEAN and compare with other available data. These data are not saved and are deleted after the user session expires.

The available data in CLEAN were reprocessed using state-of-the-art methods. While this standardized approach makes the data highly comparable and results were mostly identical to the original studies, differences with the originally used methods (e.g., aligners, genome annotations, differential expression methods, …) could result in deviations from the original findings. While we experienced no bottlenecks in downloading, processing or data storage, the manual curation of the metadata was a time-consuming task. This is illustrated by reference samples originally annotated as “untreated”, “control”, “DMSO”, “empty vector”, “timepoint 0” or simply as “sample 1”, sometimes with little to no further explanation in the manuscript and lack of source code, rendering automation of this curation step impossible. Open data initiatives like CLEAN as well as research reproducibility in general would strongly benefit from mandating of code sharing for any data processing, showing how the researcher started at *a* and ended at *z* and/or stricter guidelines for annotation of uploaded data in public repositories (11–13).

CLEAN potentiates the interactive exploration of previous studies using datasets that were generated at later time points. As we have demonstrated with the 2 use cases, this functionality can be employed to directly compare the findings of different studies or to perform GSEA using gene sets that that were published later. Recent studies, as well as our ongoing work, suggest that genomic alterations in *MYCN* and *ALK*, which are characteristic for high-risk NB, result in specific vulnerabilities to DDR inhibition (3,5). Using CLEAN we identified several drugs (i.e., lorlatinib, dactolisib and elumusertib) with different targets that resulted in 109 commonly differentially expressed genes in different NB cell lines. The identification of lorlatinib is in agreement with our recent wet lab experiments (6). Further, several of these genes have been previously investigated as putative therapeutic targets (e.g., *RRM2, CDK2, CHEK1* (*22,25*)), illustrating how CLEAN can be used to identify novel targets that can be selected for further experimental validation.

We further illustrated the strength of CLEAN in applying newly published GSEA datasets on older studies by performing a protein kinase S/T phosphosite GSEA on existing phosphoproteomics datasets, using the recently published atlas of substrate specificities for the human serine/threonine kinome (21). This analysis confirmed a previously suggested and strong compensatory activity of ATM upon ATR inhibition (3,6,27). Furthermore, a role for several Ribosome S6 Kinase (RSK) proteins was suggested in mediating the effects of both ATR and ALK inhibition.

Interestingly, RSK has been previously demonstrated to suppress ATM activity (33).

As phosphomotifs of protein kinases from similar phylogenies are highly similar, there’s a risk of mispredictions for related protein kinases. This could explain why the reduced ATR activity upon ATR inhibition was rather weak and non-significant (i.e., overlapping hyperphosphorylated ATM S/TQ sites weaken the signal) and also makes it hard to distinguish between the different RSKs based on these results.

In conclusion, CLEAN is a highly interactive and easy to use web application that centralizes all NB cell line data in a standardized format, providing a rich resource for future preclinical neuroblastoma research. We demonstrated with 2 use cases that CLEAN can be used to independently generate new hypothesis and identify novel putative drug targets. Additionally, CLEAN is also suitable to orthogonally validate findings derived from newly developed computational approaches, as an alternative to costly and time-consuming wet-lab experimental validation experiments.

## DATA AVAILABILITY STATEMENT

All results provided in this manuscript are based on previously published, publicly available data. All these data are accessible in CLEAN at https://ccgg.ugent.be/shiny/CLEAN/.

## METHODS

### Data collection

To find relevant RNA-Seq data for CLEAN, we selected all studies from the GEO (https://www.ncbi.nlm.nih.gov/geo/) (14) that fulfilled the following search criteria: “Neuroblastoma [Title]”, “Homo sapiens”, “Expression profiling by high throughput sequencing”, “Cell line”, “Sample count from 4 to 10000” (implying minimally 2 replicates for each condition and control). The selected studies were manually curated and studies not focusing on NB, with a focus on comparing cell lines to xenografts, without any form of perturbation or without biological replicates were excluded. Additionally, data from submissions with no associated, published study were also excluded. When multiple submissions were found for a single study, data were merged. Samples were organized into either baseline (control) or perturbation (condition), where all perturbations could be categorized into one of the following: compound treatment (e.g., retinoic acid), drug inhibition, chemo, KD, KO or OE. We added information about concentration and treatment duration if this information was available.

A similar strategy was followed to find relevant phosphoproteomics data in the PRX (https://www.proteomexchange.org/) (15). We applied a simple search for “neuroblastoma” and selected only studies where phosphoproteomics data were utilized, making no discrimination on mass spectrometry method. Datasets were excluded and samples organized in the same fashion as for RNA-Seq.

### Data processing

We used HISAT2 (34) to align reads from the GEO-derived fastq files to the GRCh38 reference genome. We excluded samples with an alignment rate below 75%. If this exclusion resulted in a lack of replicates, the study was excluded altogether. Subsequent quantification and annotation were performed with HTSeq (35) using GENCODE 29. For normalization of counts and differential expression analysis of RNA-Seq, we used the DESeq2 package (36). Raw phosphoproteomics intensity data were quantified using MaxQuant (37). Differential expression/phosphorylation was performed using the DEP package (38).

### Shiny web application and structure of CLEAN

The current version of CLEAN runs inside a Conda, on R (39) and and is built in Shiny (40). Tables utilize the DT package (41), plots use ggplot2 (42) and plotly (43). See Supplementary file 3 for all packages and versions used for the Conda environment. It is hosted at a webserver at Ghent University.

### Gene Set Enrichment Analysis (GSEA)

CLEAN provides both a classical GSEA (also referred to as an overrepresentation analysis) and a preranked GSEA.

The classical GSEA is performed using a one-sided Fisher’s exact test on a contingency table that compares the number of genes/proteins/phosphosites below and above the user-defined DE/DP thresholds to their presence in a specified gene set. Odds ratios (OR, indicative of the extent of the enrichment), *P* values and *P_adj_* values (multiple testing correction using the Benjamini-Hochberg method) are reported by CLEAN.

Preranked GSEA was performed using the R *fgsea* package (*fgseaMultilevel* function, default parameters) with ranking based on the DESeq2 statistic for RNA-Seq data or –log10(*P*) * Sign(log2FC) for phosphoproteomics data.

Relevant gene sets were derived from the MSigDB v2023.1 (20). For phosphoproteomics analyses, we also added protein kinases – phosphosite motif dataset. These data were derived from the supplementary information provided by Johnson et al. (21), using the top 1% predicted protein kinases for each phosphomotif and converted protein names to gene names for both kinases and target sites.

### Integrated data and links

We added links to other resources frequently used in the field: *GeneCards (*https://www.genecards.org/) for general gene information (16), *ChEMBL (*https://www.ebi.ac.uk/chembl/) for up-to-date, target-specific drug information (17), *Ensembl (*https://www.ensembl.org/) for detailed genomic information (18) and the *R2: Genomics Analysis and Visualization Platform* (http://r2.amc.nl) for Kaplan-Meier analysis on the Kocak cohort (44). We also downloaded the CRISPR/cas9 generated gene scores from the *DepMap* database (https://depmap.org/portal/download/all/, (22Q2)) and integrated them directly in CLEAN (19).

## Suppl. file descriptions

**Supplementary file 1. Tutorial and explanation of use case 1.**

A short video describing in detail how to use CLEAN, focusing on the search- and RNA-Seq-tab.

**Supplementary file 2. Tutorial and explanation of use case 2.**

A short video describing in detail how to use CLEAN, focusing on the phosphoproteomics tab.

**Supplementary file 3. Overview of packages used to run CLEAN.**

YAML (.yml) file with packages and dependencies for the Conda environment used to generate the code for CLEAN.

## ACKNOWLEDGEMENT

The authors thank all members from the Van den Eynden lab, Palmer lab and Hallberg lab for extensively testing CLEAN and providing valuable input.

This work was supported by the Ghent University Special Research Fund Starting Grant (JVdE BOF.STG.2019.0073.01), the Research Foundation Flanders (FS, JVdE, RHP FWO.OPR.2023.0063.01 and JVdE V424522N), Belgian Foundation Against Cancer (FS, JVdE STI.STK.2023.0017.02), Stand up to Cancer, the Flemish cancer society (AC STI.VLK.2022.0013.01), the Swedish Cancer Society (RHP CAN21/01549) and the Swedish Childhood Cancer Foundation (RHP 2019-0078 and JS 2021-0068).

